# StereoMate: 3D Stereological Automated Analysis of Biological Structures

**DOI:** 10.1101/648337

**Authors:** Steven J. West, Damien Bonboire, David L Bennett

## Abstract

Tissue clearing methods offer great promise to understand tissue organisation, but also present serious technical challenges. Generating high quality tissue labelling, developing tools for demonstrably reliable and accurate extraction, and eliminating baises through stereological technique, will establish a high standard for 3D quantitative data from cleared tissue. These challenges are met with StereoMate, an open-source image analysis framework for immunofluorescent labelling in cleared tissue. The platform facilitates the development of image segmentation protocols with rigorous validation, and extraction of object-level data in an automated and stereological manner. Mouse dorsal root ganglion neurones were assessed to validate this platform, which revealed a profound loss and shift in neurone size, and loss of axonal input and synaptic terminations within the spinal dorsal horn following their injury. In conclusion, the StereoMate platform provides a general-purpose automated stereological analysis platform to generate rich and unbiased object-level datasets from immunofluorescent data.

## Introduction

Recent interest in tissue clearing methods has resulted in the development of several new protocols, all promising to deliver high quality 3D reconstructions of nervous system tissue^1–9^.

A key challenge with these datasets is knowledge extraction^10^, which will be improved with well-validated automated image processing algorithms. Furthermore, any analysis framework should be capable of overcoming inherent biases when analysing 3D samples of diverse and heterogenous objects, in line with stereological principles^11,12^.

This paper describes the StereoMate platform, a pipeline consisting of tissue clearing and immunohistochemical labelling protocols and an automated stereological image analysis framework. This methodology provides protocols for deep tissue immunohistochemical labelling and imaging for 3D reconstruction of biological objects, and an analysis framework for validated knowledge extraction. This workflow is demonstrated on mouse dorsal root ganglion neurones, assessing both the cleared ganglion and its connections with the dorsal horn in naïve tissue and tissue subject to peripheral nerve injury^13^.

## Results

### The StereoMate Platform: Tissue Clearing and Immunofluroescence Labelling

A key challenge with tissue clearing methodologies is their impact on antibody affinity to their antigens^14,15^, and so the first goal was to develop a methodology that removed lipid membrane barriers to enhance antibody penetration into tissue, yet still maintained tissue antigenicity (Figure 1a).

**Figure 1:**
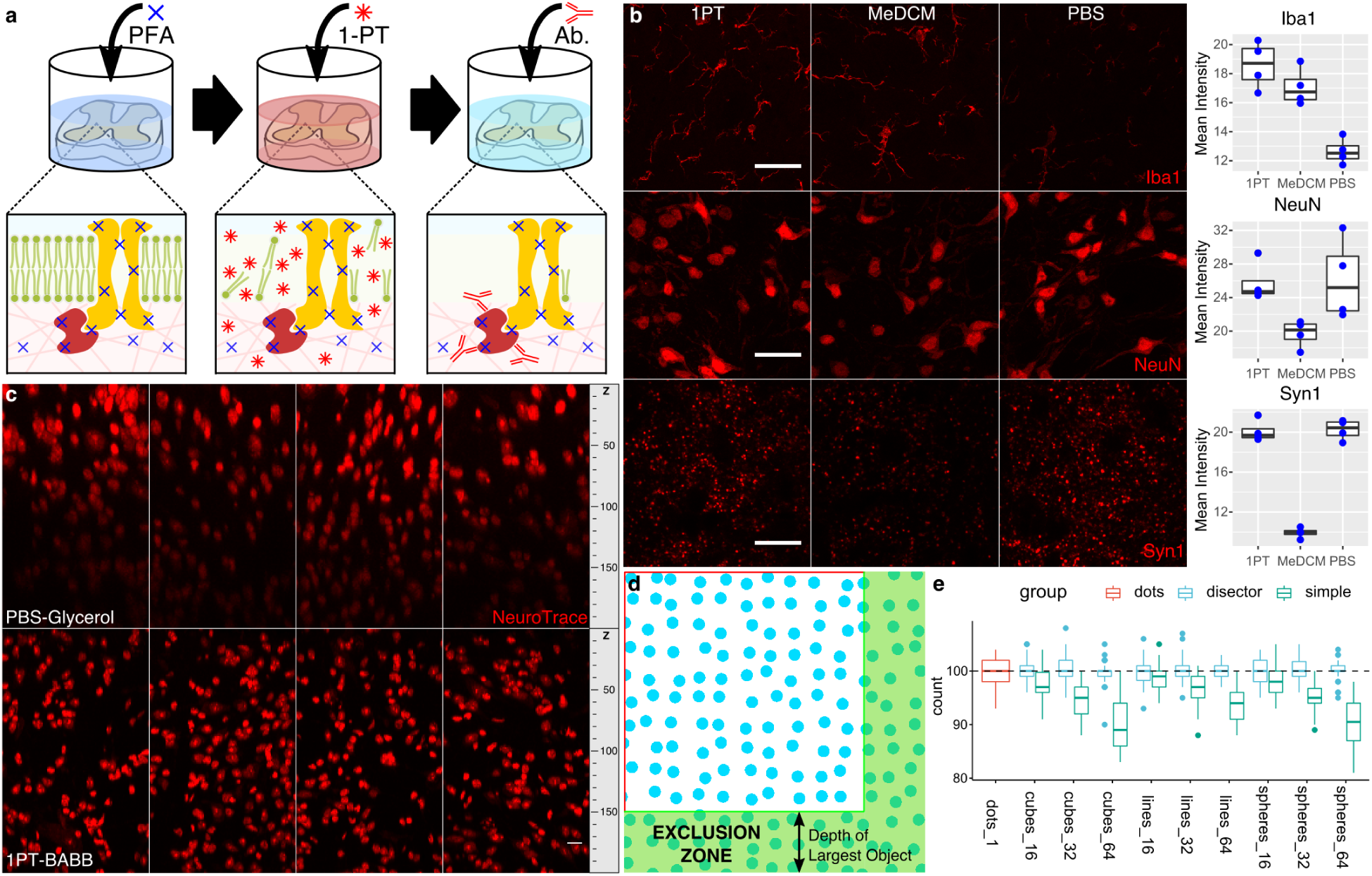
The StereoMate Histological and Image Analysis Workflow. a, Thick blocks of nervous tissue are first fixed in formaldehyde solution (PFA) to crosslink protein, then incubated in buffered 1-propanol (1-PT) to elute lipids, and finally proteins of interested are labelled using standard immunofluorescence methods (Ab.). b, Using buffered solvent to elute lipids produces the most consistent signal for a range of common antigens in dorsal horn spinal cord (scale bar: 20um for Iba1 and NeuN, 10um for Syn1). c, Dorsal horn tissue labelled with NeuroTrace viewed in YZ dimension, with X-projection applied, showing that dehydration and mounting in BABB results in good retention of fluorescence signal up to and beyond 200um Z-depth in dorsal horn compared to mounting in a glycerol-based mountant. d, The ROI Disector probe was tested with a synthetic dataset comprising systematically randomly distributed objects in 2D. This schematic illustrates an example the spheres_64 data, showing an application of the Exclusion Zone to eliminate bias in object sampling. e, Boxplots of numbers obtained from systematic random distributions of objects of different size (16, 32, 64 pixel diameter), shape (sphere, cube, line), and orientation (line: vertical, horizontal, diagonal). All distributions were sampled with an area that should contain 100 objects on average. Dots represent samples of single pixels, and show no bias. Simple counts show progressively more bias (under-counting) of larger objects. Applying the ROI DiSector corrects this bias to identify object numbers around 100. 1-PT: 1-propanol + 0.3% TEA. MeDCM: Methanol-dichloromethane. Syn1: Synapsin 1.

Different solvents, known to effectively extract lipids from biological mixtures^16,17^, were utilised for tissue lipid elution and tested for their ability to retain antigenicity across antigens from different cellular structures (including cytosolic, cytoskeletal, nuclear and membranous), in mouse spinal cord. This revealed that across all antigens, using a propanol-based lipid elution strategy retained immunofluorescence signal (Figure 1b).

A second challenge with tissue clearing preparations is optical access, which can be resolved with refractive index (RI) matching. Lipid elution results in a tissue preparation largely composed of protein, which varies in refractive index from ∼1.43 when hydrated to >1.5 when dehydrated^18,19^. As dehydrated protein closely matches the RI of borosilicate glass coverslips and immersion oil, and antibody-tagged fluorophores remain fluorescent in tissue after solvent dehydration and organic media mounting (unlike fluorescent proteins^20^), a design decision was made to exploit these features by dehydrating tissue and mounting in BABB (Figure 1a). This resulted in well matched RI throughout the light path, and thus eliminated RI variance as the Z stack was traversed (Supplementary Figure 1). This allowed confocal imaging data to be acquired at high resolution and with minimal impact on signal strength throughout the Z stack (Figure 1c).

Thus, lipid elution with propanol provides antibody access to antigens for optimal tissue labelling, and RI matching during tissue mounting gives optical access throughout tissue blocks. Together these facilitate the generation of high quality 3D reconstructions of immunofluorescently labelled tissue elements.

### The StereoMate Platform: Automated and Unbiased Image Analysis

To effectively extract object-level knowledge from the resulting 3D datasets in an unbiased and demonstrably reliable manner, the StereoMate platform was developed. The workflow involves delineating regions of interest (ROIs) for object analysis (*ROI Manager plugin*), interactively establishing an image segmentation procedure to extract biological objects of interest from the raw image data (*Threshold Manager plugin*), and demonstrating the accuracy of this segmentation procedure through assessing a sample of objects (*Object Manager plugin*). Finally, the image segmentation protocol is used to derive the population of biological objects for automatic assessment in an ROI-dependent manner, using a novel stereological probe (*StereoMate Analysis plugin*).

User-defined ROIs can be optionally defined on input images with the ROI Manager Plugin (Supplementary Figure 2), assigning biological objects to regions within the tissue. The StereoMate framework also optionally provides an Image Deconvolution plugin, with built-in PSFs, to use prior to image thresholding (*StereoMate Deconvolution plugin*), which improves image resolution and signal contrast with high resolution imaging (Supplementary Figure 3).

The Threshold Manager Plugin plays the crucial role of defining a segmentation procedure. This is developed interactively by a user, by assembling a sequence of image processing operations. These operations act on the current image, and the user can initially decide through visual inspection when a suitably accurate segmentation operation is defined (Supplementary Figure 4).

The accuracy of object segmentation underpins the quality of the object-level data extracted in the analysis step. It is therefore essential to assess, on a sample of segmented objects, the accuracy of this segmentation. The Object Manager Plugin facilitates this crucial and necessary step. The plugin displays the raw pixel data overlaid with an image segmentation procedure defined by the Threshold Manager. The user can generate sample selections of segmented objects to inspect their 3D characteristics and classify them (Supplementary Figure 5).

Common segmentation problems include inaccurate object representations that do not fit to the underlying pixels, and the connection of independent objects during segmentation. These problems are overcome through sequentially optimising and assessing the segmentation procedure with the Threshold and Object Manager plugins.

The Object Manager further provides the ability to develop an object filter to define a sub-group of biological objects for analysis. The object filter can be either a simple high- and low- pass filter on a single feature of the objects, or a complex machine learning classifier, applied to multiple features, trained by the user’s classification of a sample of objects (Supplementary Figure 5).

Once object segmentation has been optimised, object-level data is extracted with the StereoMate Analysis Plugin. The assessment of objects from a tissue segment is subject to a range of biases, which can be eliminated through the use of stereological techniques (Supplementary Figure 6 & Figure 1d). To overcome this bias in biologically relevant regions, the ROI Di-Sector was developed.

This di-sector probe applies the logic of the optical di-sector^21,22^ to user defined ROIs of a 3D sample (Supplementary Figure 7). The algorithm ensures an unbasied set of fully reconstructed objects are returned from each ROI, by replacing acceptance lines with exclusion zones on the image boundaries of sample z stacks (Figure 1d & 1e). Each fully reconstructed object in the unbiased sample is measured for size, shape, location and intensity data.

The StereoMate Workflow provides robust histological labelling of thick tissue blocks for detailed reconstruction of labelled elements, and an image analysis workflow that delivers demonstrably accurate, reliable and comprehensive object-level quantitative data for statistical analysis.

This method has been applied to both mouse wholemount dorsal root ganglia (DRG) preparations to assess total neurone numbers in DRG, and to dorsal horn to assess DRG axonal input and synaptic puncta, presented next.

### Assessment of neurones in wholemount DRG

A key advantage of tissue clearing methods is their ability to facilitate large scale 3D reconstructions of biological tissues. The ability to reconstruct entire regions of tissue would enable the entire population of biological objects within it to be analysed. This negates the need for any stereological estimates and offers the most comprehensive assessment. To achieve this, the relatively small dorsal root ganglion (DRG) was chosen for labelling and assessment using the StereoMate workflow.

As DRG neurones are densely packed, a strategy was devised to label neuronal nuclei, which are more easily resolved than neuronal soma labelling. TDP43 was identified from single cell transcriptomic data from DRG^23^ as a highly expressed nuclear antigen across all DRG neurone sub-types.

Labelling of TDP43 in wholemount DRGs revealed strong expression and localisation to neuronal nuclei (Figure 2a, Supplementary Video 1), but also within satellite glial and schwann cell nuclei, consistent with its expression globally in all healthy nuclei^24^. The nuclear labelling was well maintained both to the centre of the tissue, indicating sufficient penetration of antibody, and to the bottom of the Z stack, demonstrating good matching of refractive index throughout the light path (see Supplementary Video 1).

**Figure 2:**
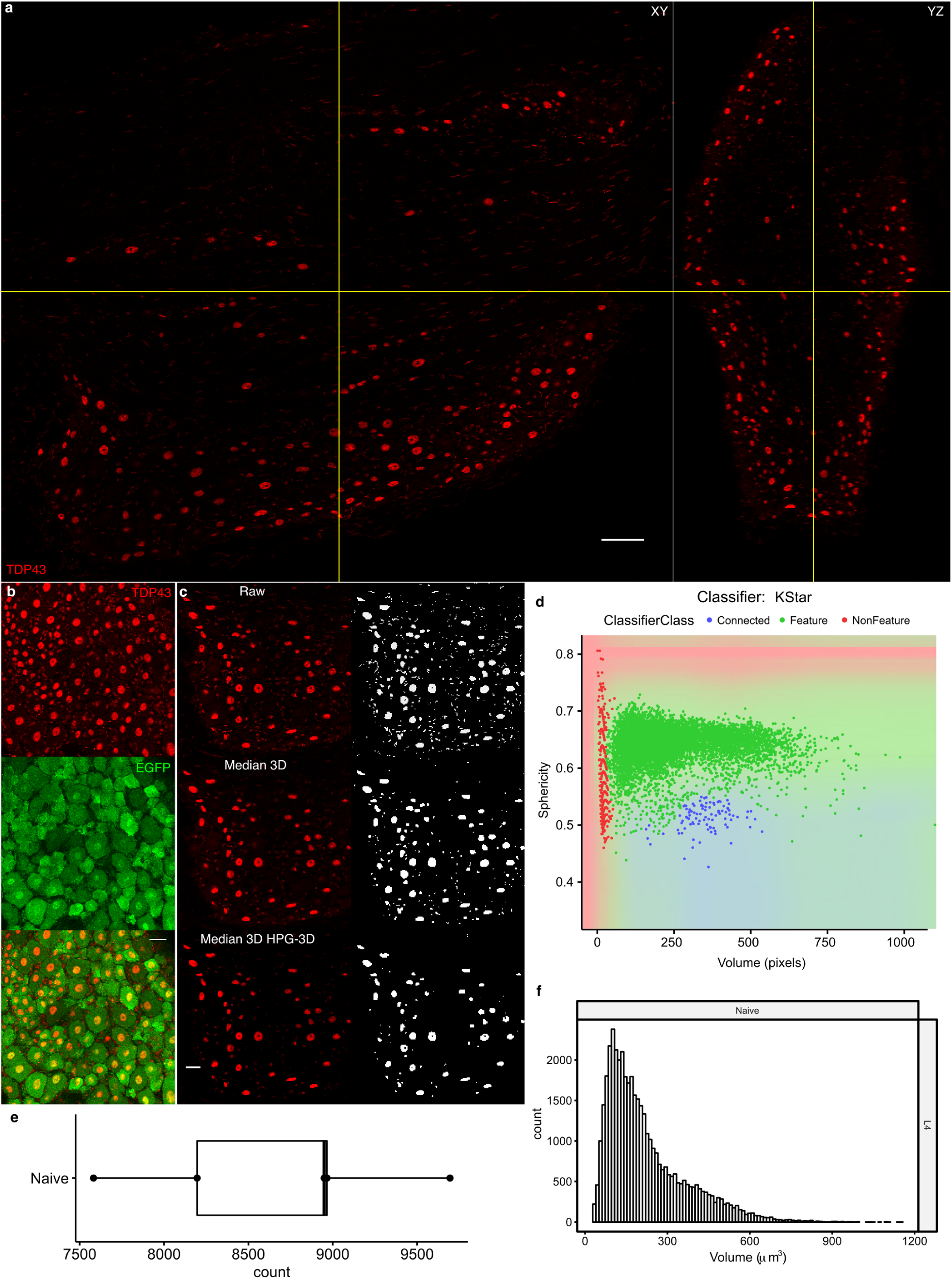
Assessment of wholemount DRG neurones. a, Orthogonal Views of a cleared DRG labelled with TDP43 antibody, showing the XY and YZ views of the tissue block. Note the even labelling of TDP43+ nuclei to the centre and bottom (in Z) of the tissue block. b, Panel showing TDP43 and EGFP labelling in DRG indicating that TDP43 is expressed in all DRG neuronal nuclei. c, Overview of image filters applied prior to image segmentation. With the application of a low-pass filter (Median 3D) and a high-pass filter (High-Pass Gaussian filter 3d), it is possible to repress un-desired high- and low- frequency components, and therefore focus on extracting the desired neuronal nuclei, improving segmented object data quality. d, Scatterplot of segmented DRG objects showing the data distribution by size and sphericity measures. Each object has been classified by a KStar machine learning classifier as FEATURE (neuronal nucleus, green), NON-FEATURE (glial nucleus, red) or CONNECTED (two or more neuronal nuclei connected together, blue) The vast majority of objects are neuronal nuclei, due to image filtering. The decision boundary of the KStar classifier colours the scatterplot below the datapoints. e, Boxplot of DRG neuronal nuclear number, showing 8,677.6±809.9 (mean±SD; n=5) DRG neurones. f, Histogram of DRG neuronal nuclear size of all DRGs assessed (n=5). A multi-modal distribution of neuronal nuclear size can be seen.

To confirm TDP43 labelled all neuronal nuclei, a transgenic mouse where EGFP was expressed under the Advillin promotor (known to be expressed across the range of DRG neurones^25,26^) were co-labelled with anti-TDP43 and anti-EGFP antibodies (Figure 2b, Supplementary Video 2). In all tissue analysed, neuronal cell bodies showed varying levels of EGFP expression, and each EGFP+ neurone possessed a TDP43+ nucleus.

A TDP43+ nuclear object segmentation protocol was established with the Threshold Manager plugin and validated with the Object Manager plugin. It was found that optimal segmentation was achieved with a narrow confocal pinhole (0.5AU), application of a bandpass filter to suppress non-neuronal signal, and segmenting the histogram by fitting it to a multi-modal model (Figure 2c).

To separate the remaining glial nuclei from neuronal nuclei, a machine learning classifier was trained from a sample of user-classified TDP43+ segmented nuclei (Figure 2D). This classifier showed robust performance (∼99% accuracy) on a large independent test set (n=990, accuracy 99.5%, Supplementary Figure 8), providing high confidence in the quality of the classifier. The StereoMate Analyser plugin was then used to extract all objects classified as neuronal nuclei for analysis.

To explore the data, some basic plots were generated (Figure 2e-f). Adult mouse L4 DRGs were found to contain 8,677.6±809.9 (mean±SD; n=5) DRG neurones. A plot of the size histogram of DRG nuclei reveals a novel multi-modal distribution with at least three major divisions. This extends previous 2D analyses of DRG soma size, which revealed a bimodal distribution^27^.

### StereoMate Compared to Manual Stereological Assessment

To enable a direct comparison between the StereoMate analysis and a manual stereological assessment, confocal image stacks from a set of DRGs were digitally re-sampled to generate a set of systematic randomly sampled images (Figure 3a & 3b).

**Figure 3:**
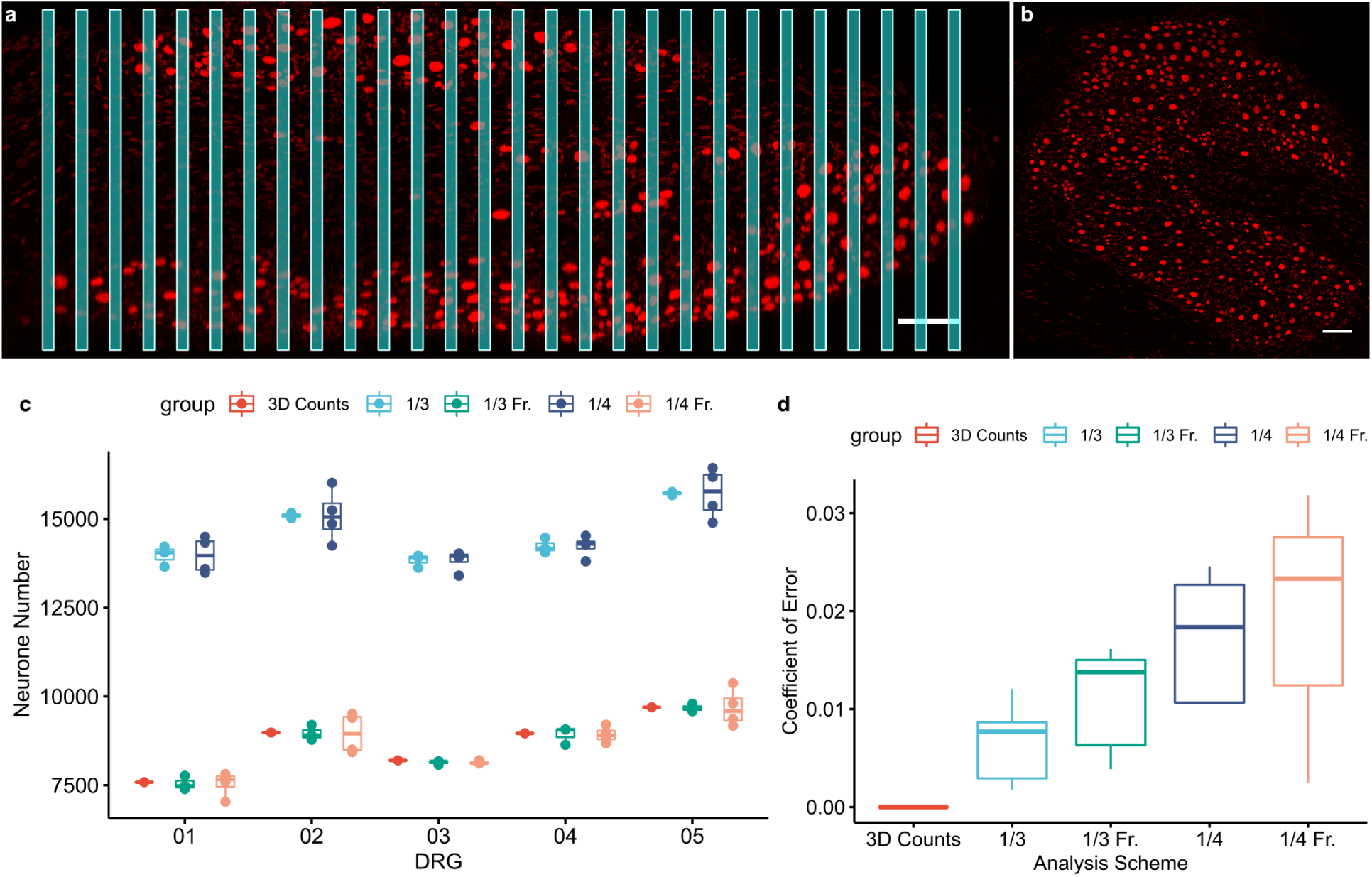
Comparison of StereoMate analysis to Manual Stereological analysis. a, Confocal image stacks of DRGs were re-sampled for manual stereological analysis using a systematic random sampling scheme. This schematic illustrates the sampling of one DRG to obtain a 1/3 fraction of the total DRG, according to the fractionator sampling rules. b, a single sample derived from a, projected to show the typical TDP43 signal to be assessed stereologically. c, Boxplot showing the 3D counts, simple nuclear counts with 3x 1/3 and 4× 1/4 sampling (1/3 SC and 1/4 SC, respectively), and optical disector (fractionator) counts with 3x 1/3 and 4x 1/4 sampling (1/3 Fr. and 1/4 Fr. respectively). Simple counts over-estimate DRG neuronal number as predicted by stereological theory. Stereological analysis brings the estimate close to 3D object counts. d, The Coefficient of Error was measured for both simple and stereological counts. Quantification of whole DRGs (3D Counts) has no sampling error, as indicated. Simple and stereological counts have different sampling errors depending on the sampling and analysis scheme. Note, simple counts under-estimate the CE, due to an over-estimate of DRG neurone number without an equivalent increase in estimate error.

Both simple counts of neuronal nuclei and the Optical Fractionator stereological method were performed on the samples, at two sampling rates (1/3 and 1/4 of the total DRG volume, Figure 3C). Across all DRGs, using simple counts resulted in a large over-estimate of neuronal numbers, a reflection of the fact that each nucleus is cut into multiple profiles and so will be ‘double counted’ using this methodology^11,12^. However, application of stereological technique corrected this to bring the estimate of neuronal numbers in each DRG to cluster around the counts observed from 3D reconstruction, thus validating both the 3D reconstruction numbers and the stereological methodology.

To assess the degree of noise added by the different analysis schemes, the coefficient of error (CE) was measured. The CE gives a measure of variation independent of the sample size and sample mean. Figure 3D shows a boxplot of the CE measured across all assessed DRGs, showing that CE tends to increase with reduced sampling. Interestingly, the CE is higher on average for stereological assessment than with simple counts, due to simple counts over-estimating the object number, but not increasing the variance by the same degree, which artifically brings down the CE. Importantly, whereas each sampling scheme shows an inherent variation due to sampling, the data from fully reconstructed DRGs has no sampling error, and thus leads to more powerful and reliable data.

Comparisons were made in the time taken to assess each DRG. Manual stereological assessment at 1/3 sampling took approximately 3-4 hours per DRG, compared to ∼2 minutes when running the StereoMate Analysis plugin, optimised and validated to assess all DRG neurones.

Finally, whereas manual stereological analysis only yields a single estimate of object number, the StereoMate analysis platform yields an unbiased sample of objects which can be measured in various ways. This multivariate dataset allows a much wider variety of analyses to be performed across different object measures and their derivatives.

In conclusion, the StereoMate framework overcomes subtle biases present in manual stereological analysis, and delivers a powerful and rich dataset for detailed assessment of biological structure.

### TDP43+ Neuronal Nuclei are lost and shift size distribution following nerve injury

To test the hypothesis that peripheral nerve injury results in a loss of TDP43+ neuronal nuclei, whole DRGs from 6 week post spared-nerve injury^28^ (ipsilateral [L4i] and contralateral [L4c] L4 DRGs) and sham surgery (ipsilateral L4 DRGs only) were cleared and labelled for TDP43 (Figure 4a).

**Figure 4:**
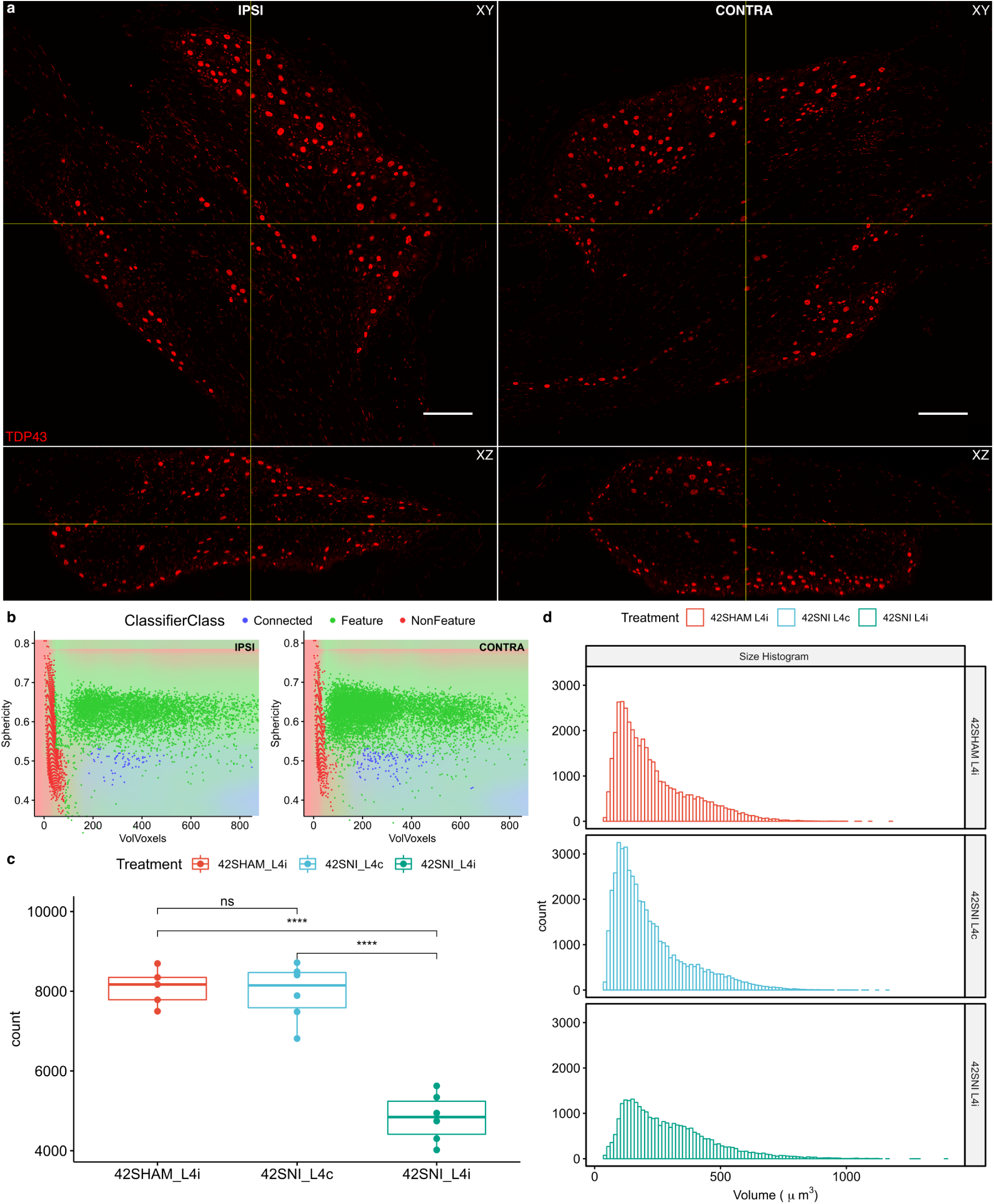
DRG neurones are lost following peripheral nerve injury. a, Example orthogonal views of DRGs labelled with TDP43 anti-antibody. The two DRGs are from the same sample, from the ipsilateral and contralateral L4 DRG. b, Scatterplot of segmented DRG objects from DRG images in (a), showing the data distribution by size and sphericity measures. Each object has been classified by a newly built KStar machine learning classifier as FEATURE (neuronal nucleus, green), NON-FEATURE (glial nucleus, red) or CONNECTED (two or more neuronal nuclei connected together, blue) The decision boundary of the KStar classifier colours the scatterplot below the datapoints. The cloud of FEATURE objects (neuronal nuclei, green) in the ipsilateral data appears to be significantly less than in the contralateral data. c, Boxplots of total DRG neuronal numbers for SNI L4i, L4c and SHAM L4i groups. Data compared using a t.test with adjusted p-values for multiple comparisons using the Holm method (****, p<0.0001, n=6 for SNI groups, n=5 for SHAM group). e, Size histograms of summed DRG data, revealing a trimodal distribution for SNI L4c and SHAM L4i, but a large reduction in histogram size, and predominant loss of small DRG neurone populations in SNI L4i group.

A segmentation procedure was developed with the Threshold Manager plugin identical to the previous procedure for naive tissue. Again optimal segmentation was achieved with a narrow confocal pinhole (0.5AU), application of a bandpass filter to suppress non-neuronal signal, and segmenting the image by fitting the histogram to a multi-modal model.

A new machine learning classifier for this second dataset was trained from a sample of user-classified TDP43+ nuclei via the Object Manager plugin (Figure 4b). This classifier showed robust performance on a large independent test set (n=1,000, accuracy 99.5%), providing high confidence in the quality of the new classifier. The StereoMate Analyser plugin was then used to extract all objects classified as neuronal nuclei for analysis.

Comparisons of total DRG neurone number between groups revealed a significant and profound reduction in TDP43+ neurone number in the SNI injured DRGs (Figure 4c). An approximately 3/8 reduction in TDP43+ neurone number is seen at this 6 week time point post nerve injury.

To identify which population(s) of neurones are affected at this timepoint, the size histograms were compared between groups. This revealed a significant reduction in the first two peaks of the size histogram, revealing an impact on small C fibres (Figure 4d).

### Assessment of primary afferent axons and synapses in Dorsal Horn

The complete reconstruction of the DRG reveals the power of analysing the entire population of biological objects within a complete region of tissue. However, this does not utilise the 3D stereological di-sector previously outlined (Figure 1D). To employ the 3D stereological di-sector, and to further explore DRG neurone anatomy, adult mouse dorsal horn spinal cord tissue was dissected and prepared for tissue clearing and labelling.

Utilising the Advillin-EGFP transgenic mouse, EGFP+ primary afferent axons were labelled and the distribution of axonal fragments in dorsal horn samples were assessed. Furthermore, two pre-synaptic terminal markers were assessed: Synapsin 1 (Syn1), and Synaptoporin (SynPor.).

As the dorsal horn forms a continuous stretch of grey matter throughout the length of the spinal cord, it is impractical and unnecessary to reconstruct the entire structure. To avoid bias when assessing a set of fully reconstructed objects within this sample (as exemplified Figure 1d & 1e), a stereological di-sector probe must be deployed. The ROI Di-Sector was therefore utilised for assessing the pre-synaptic terminal labels.

Likewise, it can be impractical and unnecessary to reconstruct entire objects. Where whole object reconstruction is not practical, it is possible to simply assess the fragments of objects that exist within a sampled region, which will not be subject to biases as when reconstructing whole objects. This was employed for assessing the EGFP+ primary afferent axonal imput to the dorsal horn, which are otherwise impractical to reconstruct entirely. However, it is always more desirable to reconstruct entire objects and regions where practical, as these offer the most comprehensive datasets.

Examples of the dorsal horn sampling and image processing are shown in Figure 5a & 5b. Confocal image stacks were taken from the mid-region of the dorsal horn, overlying laminae I-III. Deconvolution was used to improve object constrast and reduce signal blur in all datasets. Both EGFP and pre-synaptic terminal signals were effectively segmented with the Threshold Manager plugin post deconvolution by using a 3D median filter and fitting a multi-modal model to the histogram (Figure 5b).

**Figure 5:**
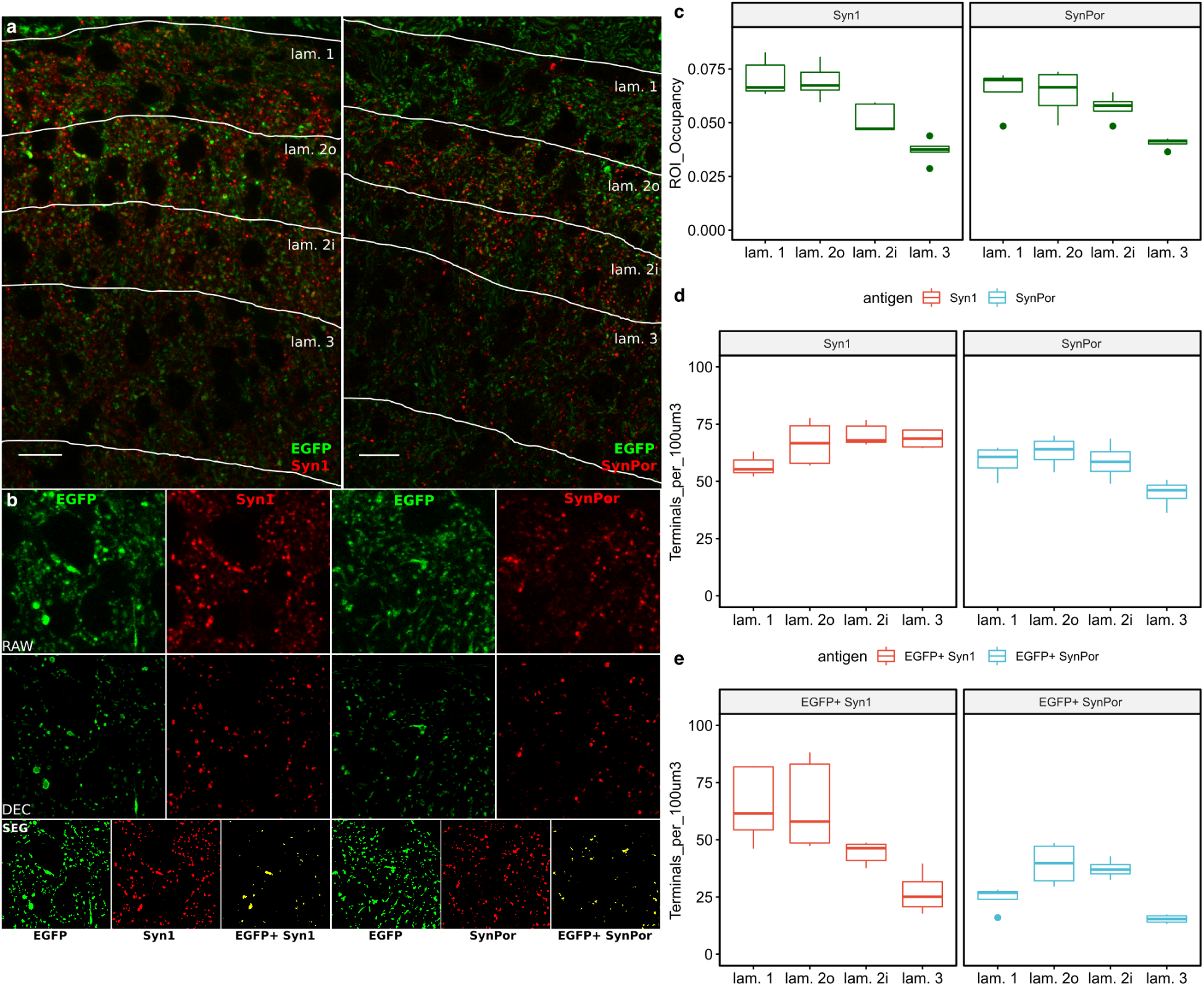
Primary afferent axonal input and pre-synaptic terminal assessment. a, Confocal Z slice of mid superficial dorsal horn, showing EGFP, Syn1 and SynPor labelling. Laminae are delineated and labelled. b, Image processing of EGFP and pre-synaptic terminal markers. Example of raw data (RAW), deconvolved images (DEC), and segmented images (SEG). The Segmented images show the three labels quantitatively assessed in (c-e): EGFP signal, Pre-synaptic terminal signal (Syn1 or SynPor), and EGFP+ Pre-synaptic terminal signal (EGFP+ & Syn1+ or SynPor+). c, ROI Occupancy of EGFP axon signal per lamina of the dorsal horn, for both Syn1 and SynPor datasets. d, Total pre-synaptic terminal density per lamina of the dorsal horn, for both Syn1 and SynPor labelling. e, EGFP+ Pre-synaptic terminal density per lamina of the dorsal horn, for both Syn1 and SynPor labelling.

The Object Manager plugin was used to assess the EGFP and pre-synaptic terminal segmentation quality. 200 objects were assessed for each dataset, and a good correspondence was observed between the segmented objects and the underlying data they were derived from.

To assess the segmented objects in each lamina, the ROI Manager plugin was used to delineate laminar borders, guided by co-staining with IB4, a marker of lamina II (Figure 5a).

EGFP+ axonal input was measured as a ratio of ROI Occupancy, and showed elevated levels in superficial laminae (I & II outer), with reduced input in deeper laminae (Figure 5c), consistent across the Syn1 and SynPor datasets.

Unbiased samples of total Syn1 and SynPor puncta showed different distributions. Whereas Syn1 puncta show an increasing puncta density through to deeper laminae, synaptoporin showed its highest density in lamina II (Figure 5d).

Finally, the density of pre-synaptic puncta that reside within EGFP+ axons were assessed (Figure 5b & 5e). EGFP+ Syn1 puncta density reveals that the majority of Syn1 puncta in superficial laminae are from primary afferents, with a reduction in primary afferent pre-synaptic terminals in deeper laminae. EGFP+ SynPor puncta density shows that SynaptoPorin is enriched in primary afferent terminals in lamina II.

### EGFP+ Axons and Synaptoporin puncta are lost following peripheral nerve injury

To understand the impact of peripheral nerve injury on injured primary afferent terminals, lumbar spinal cord from 6 week post spared-nerve injury^28^ and sham surgery was cleared and labelled for EGFP and Synaptoporin.

Data was deconvolved and segmented, with typical examples of segmentataion shown in Figure 6a. Following segmentation, a random sample of 200 objects was assessed with the Object Manager, revealing a good correspondence between individual objects manually observed in the raw data with each thresholded objects in the segmented data.

**Figure 6:**
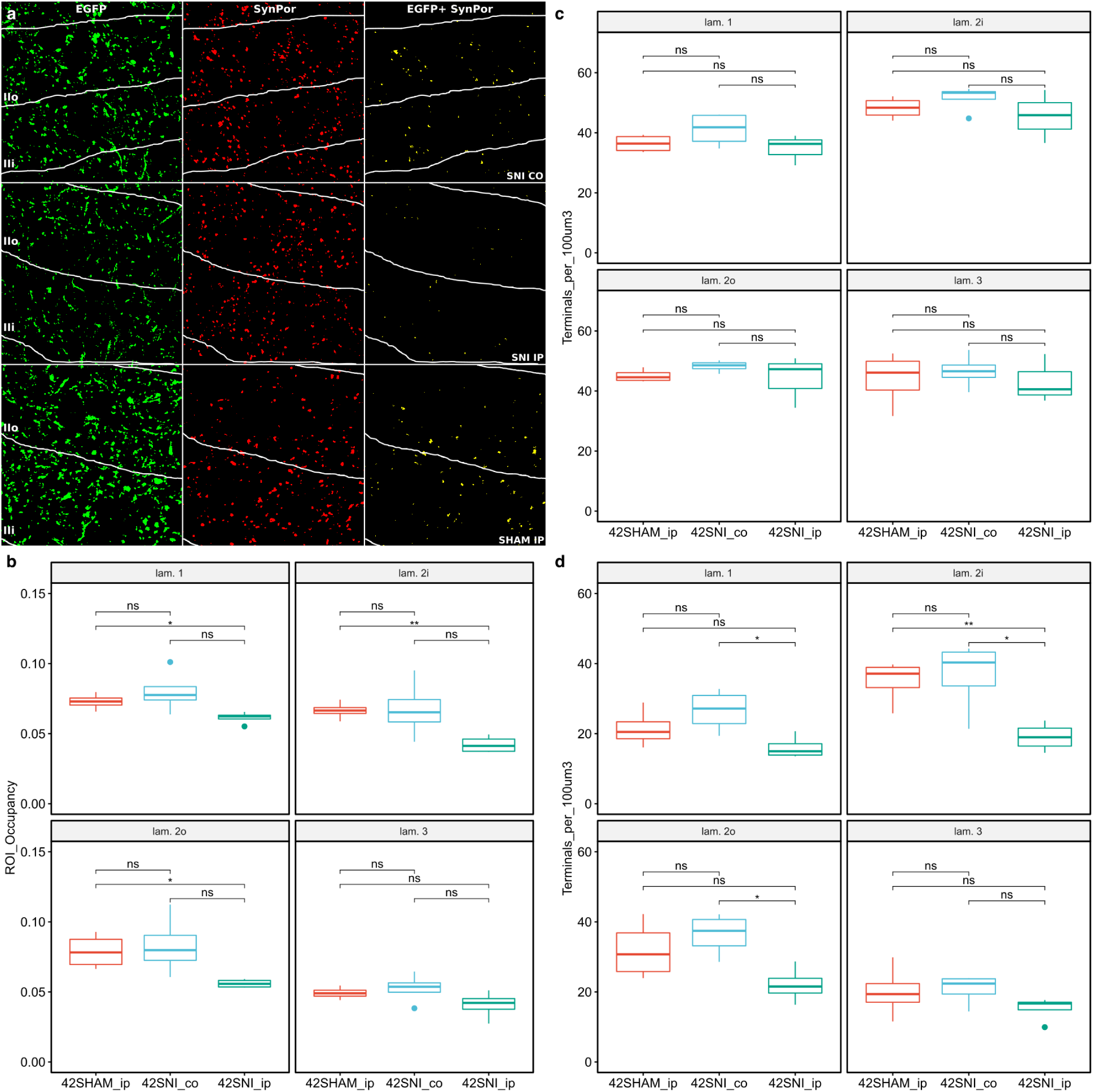
Primary afferent axons and pre-synaptic terminals are lost following peripheral nerve injury. a, Confocal Z slice of mid superficial dorsal horn (laminae 1-3), showing segmented EGFP, Syn1 and EGFP+ Pre-synaptic terminal signal (EGFP+ & Syn1+) labelling. Laminae are delineated and labelled. b, ROI Occupancy of EGFP axon signal per lamina of the dorsal horn shows a significant reduction across lamina 1-2 in 42SNI_ip relative to 42SHAM_ip. c, Total pre-synaptic terminal density per lamina of the dorsal horn shows no significant differences across all groups. d, EGFP+ & Syn1+ density per lamina of the dorsal horn reveals a significant reduction across lamina 1-2 in 42SNI_ip relative to 42SHAM_ip and 42SNI_co. Data compared using a t.test with adjusted p-values for multiple comparisons using the Holm method (*, p<0.05; **, p<0.01, n=3 for SNI groups, n=3 for SHAM group).

EGFP signal was assessed for ROI occupancy as described previously. This revealed a significant loss in EGFP fluroescence in lamina I and II between SHAM and SNI ipsilaterally, but no significant differences between SNI ipsilateral and contralateral (Figure 6b). Overall, this data suggests there is a small reduction in EGFP+ axonal input following nerve injury.

Total synaptoporin puncta numbers were assessed per lamina, which revealed no significant differences between all groups, suggesting overall synaptoporin puncta were retained following SNI injury (Figure 6c). However, assessment of synaptoporin puncta that resided within EGFP+ signal revealed some significant loss across lamina I and II, with the most pronounced in lamina II inner (Figure 6d). This suggests that the small EGFP axonal loss also corresponds loss of priamry afferent pre-synaptic terminal regions, and reflects a withdrawal of primary afferent synaptic contacts within lamina II following peripheral nerve injury.

## Discussion

This work has developed and validated a new tissue clearing and image analysis platform, optimised for use with immunohistochemical interrogation of biological structures.

The tissue clearing workflow offers improved antigenicity for a range of antigens tested, and delivers consistent high resolution imaging deep into tissue blocks with oil-immersion lenses, allowing detailed 3D reconstructions of immunofluorescently labelled structures. By using longer carbon chained solvents, which are routinely used for lipid extraction from tissues^16,17^, antigenicity is well retained. This may be due to the higher lipophilicity allowing good lipid extraction while mitigating any impact on epitopes due to excessive dehydration.

The StereoMate platform allows the user to design an image segmentation procedure interactively. This allows the user to optimise the object segmentation procedure to the current problem, but importantly enables this process to be quantitatively assessed on a sample of objects using the Object Manager plugin. The user can further develop a machine learning classifier to isolate a subset of objects based on their characteristics, as demonstrated with the DRG dataset.

The StereoMate platform can perform analyses on complete objects from whole regions, complete objects from sampled regions, and object fragments, as demonstrated with the DRG nuclei, Synaptic puncta, and EGFP+ axonal datasets, respectively. Where regional sampling is performed, whole object corrections via the ROI DiSector probe provide an unbiased set of objects for measurement. The result from any analysis is a comprehensive dataset that provides unbiased insights into the properties of objects in regions of tissue.

The StereoMate platform was applied to DRG neurones, to assess their number and properties in the DRG, and their axonal input and connections in the dorsal horn. DRGs were found to contain ∼8,000, consistent with previous stereological estimates^27^. Furthermore, a multi-modal distribution of DRG neuronal nuclear size, contrary to previous reports of 2D DRG neuronal soma distribution^29^. In dorsal horn, there is a greater axonal and synaptic input in superficial laminae from DRG neurones.

Following nerve injury, a profound loss of TDP43+ nuclei in DRG and a loss of DRG neurone axons and synaptic terminals in dorsal horn was observed. The loss of TDP43+ nuclei may reflect either loss of these neurones, or a down-regulation or re-distribution of this protein, as this is known to occur in neurodegenerative states^24^.

However, the TDP43+ nuclear loss is from smaller DRG neuronal nuclei, and DRG axonal and synaptic loss is further demonstrated in the superficial laminae (especially lamina II), where these small fibres are known to terminate^30^, which provides evidence for a functional impact between these neurones and their connection with the dorsal horn.

These results may reflect change to the IB4-binding DRG neurones. These neurones are of small size and principally form connections in lamina II of the dorsal horn^31^. These fibres are known to lose their ability to bind IB4, and their terminals show a delayed loss after injury^32,33^.

In conclusion, the StereoMate platform provides a demonstrably reliable and accurate extraction of object-level data from 3D image stacks in an automated and stereological manner. The demonstration of reliability through inspection and classification of samples of segmented objects provides confidence in the data, and produces high quality object-level measures. The richness of the datasets that are generated open up new avenues for exploring and assessing biological structures, both to address specific questions with a high degree of accuracy, but also, through exploratory data analysis, to generate further hypotheses concerning tissue structure and function.

## Acknowledgements

We would like to acknowledge the Wellcome Trust and the Strategic Award “Defining pain circuitry in health and disease” for funding this research. DLB is a senior Wellcome fellow in clinical science.

## Author Contributions

S.J.W designed all histological experiments, and conducted the DRG clearing experiments, including the manual stereological assessment. D.B conducted the spinal cord clearing experiments. S.J.W and D.B analysed the experiments with the StereoMate analysis framework. All data presentations were formatted by S.J.W. S.J.W and D.L.B wrote the manuscript.

## Online Methods

### Animals

Mice derived from the Advillin-eGFP line from GENSAT were used in this study. All animal experiments were performed in accordance with the European Community directive 86/609/EC and the UK Animals (Scientific Procedures) Act 1986, under appropriate UK Home Office personal, project and establishment licences.

### Surgery, Perfusion, Dissection

Adult animals (16-20 weeks old) were randomly allocated to receive SNI (4M/4F) or SHAM (3M/3F) surgery using aseptic technique, as previously described^28^. Briefly, animals were anaesthetised and the left sciatic nerve exposed. The common peroneal and tibial branches were ligated and transected (SNI), or nerve exposure with no manipulation (SHAM) was performed, and the surgical site sealed with sutures. Animals received 50mg/kg remifentanyl as post-surgical analgesic and were routinely monitored for 6 weeks post surgery.

Animals were terminally anaesthetised and caridally perfused with PBS followed by freshly prepared 4% PFA, and the lumbar enlargement of the spinal cord and ipsilateral and contralateral whole L4 DRGs were carefully dissected, ensuring to retain both the dorsal and spinal root. Tissue was post-fixed for 24 hours at room temperature in the same fixitive.

Lumbar spinal cord tissue was incubated in 30% sucrose in PBS for 2-3 days, embedded into OCT compound and frozen for cryostat sectioning. Lumbar spinal cord tissue was sectioned at 50um thickness, with the contralateral ventral horn marked for orientation, and sections washed in PBS for free floating immunohistochemistry. Sectioned spinal cord and whole DRG tissue was washed in PBS and stored at −80 in CI-VM1 (35% DMSO 35% Ethylene Glycol in PBS) until labelling.

### Clearing and Immunohistochemistry

Tissue clearing was performed by dehydrating in a gradient of 1-propanol containing 0.3% tirethylamine (30, 50, 75, 90, 95, 100, 100%), and washed in this solution at 37C for 24 hours. Tissue was rehydrated into PBS, and antibody labelling was performed at 37C for 7 days. Tissue was washed for 24 hours in buffer, and incubated in secondary antibody at 37C for 7 days. Tissue was washed for 24 hours in buffer, dehydrated in increasing concentrations of 1-propanol containing 0.3% triethylamine, and mounted in BABB (benzyl alcohol, benzyl benzoate, 1:2 ratio) containing 0.3% triethylamine on glass slides with rubber silicone spacers. Tissue was secured with a No 0 coverslip for confocal imaging.

### TABLE of antibodies

DRG: with anti-TDP43 (ab133547, abcam) 1:100 dilution anti-rabiit Cy3 conjugated secondary antibody (711-165-152, JIR) 1:500 dilution IB4

### Confocal Microscopy

Confocal imaging was performed on a Zeiss LSM 700 confocal microscope. The Zeiss LD LCI Plan-Apochromat 63x/1.2 Imm Corr DIC objective was used for imaging of wholemount DRG, with the correction collar set to oil and zeiss objective oil used for immersion.

Images were acquired at 1um resolution in XYZ. The laser and gain was adjusted to give optimal signal throughout the Z stack, with a gradient set to increase laser and gain through Z as appropriate for consistent signal. A tile scan and Z stack was taken to capture the entire DRG within one image.

For high resolution imaging of axonal and synaptic structures in spinal cord, the Zeiss Objective Plan-Apochromat 63x/1.4 Oil DIC objective was used. Images were acquired at 0.04×0.04×0.1um voxel size, below the Nyquist sampling rate for this objective, for deconvolution. The laser and gain was adjusted to give optimal signal throughout the Z stack, with a gradient set to increase laser and gain through Z as appropriate for consistent signal. A tile scan was taken to capture a field of view covering laminae I-III of the dorsal horn. A single z slice containing IB4+ morphological information was taken for delineating each lamina during analysis. A Z stack (8um thick) was taken of axonal and synaptic antigens for 3D stereological analysis.

### Image Analysis

The Open Source StereoMate ImageJ plugin suite was used for assessing wholemount DRG confocal image stacks and is freely available on sourceforge (), with source code on github (https://github.com/stevenjwest/StereoMate).

StereoMate consists of several plugins which sequentially process a set of images to enhance and filter the image signal, define regions of interest for object assessment, develop and verify an image segmentation strategy to derive a set of structures for assessment, and finally assess these structures in a stereological manner. Full documentation on the use of these plugins is provided on the github page for the StereoMate suite, here a step by step guide is given for the processing performed in this work.

### StereoMate Deconvolution Plugin

Axonal and synaptic confocal image stacks were deconvolved using the StereoMate Deconvolution plugin. This uses the WPL deconvolution algorithm in the Parallel Iterative Deconvolution plugin by Piotr Wendykier^34^, but also provides theoretical PSFs for a range of objectives and fluorophores. The parent directory containing all Z stacks was selected for input, and each channel was deconvolved with the appropriate PSF with Max. Iterations set to 40. The deconvolved images were automatically saved to the user-designated output directory for further analysis.

### ROI Manager Plugin

The ROI Manager plugin was use to define each lamina for analysis of synaptic and axonal structures. The parent directory containing all images was selected for input, each image containing IB4+ morphological information was opened, and each ROI corresponding to a lamina of the dorsal horn was interactively defined. The ROIs were automatically saved to the user-designated output directory for use in further analysis.

### Threshold Manager Plugin

The Threshold Manager plugin was used to define the image segmentation protocol for each dataset. This plugin provides an interactive environment for the user to define and inspect an image segmentation procedure, consisting of image filtering and auto-thresholding algorithms. Any imagej plugin can be used within this interactive plugin for processing the image, and a number of common procedures are defined in the drop-down menus.

The parent directory containing all Z stacks was selected for input, and the bit depth was set to 8. The segmentation protocols images were saved to the user-designated output directory for use in further analysis.

The optimal DRG segmentation procedure, as confirmed by the Object Manager plugin, used a Median 3D filter (radius 1) as a low-pass filter, followed by a plugin which blurred the image with a 3D Gaussian filter (radius 10) and subtracted this blurred image from the original image (the built-in High-Pass Gaussian filter), which acts as a high-pass filter. The DRG image was segmented using the built-in Multi-OTSU segmentation algorithm, using 5 levels and defining the background as level 1.

The optimal segmentation procedure of the deconvolved image data for both axons and synaptic puncta used a Median 3D filter (radius 1) as a low-pass filter, followed by segmentation with the built-in Multi-OTSU segmentation algorithm, using 5 levels and defining the background as level 0.

### Object Manager Plugin

The Object Manager plugin was used to explore the segmented data and confirm its accuracy with random or biased selections of objects. This plugin also allows the user to define simple object filters to select objects based on a simple high- and low- pass value for one attribute. It further allows the user to build a machine learning classifier to define object classes based on multiple attributes, using the Weka Workbench library^35^.

The parent directory containing all Z stacks was selected for input. The relevant image segmentation procedure file was selected from the output of the Threshold Manager plugin, and all objects were assessed with 6-connected Object Connectivity. TDP43 labelling in DRG, and pre-synaptic terminal labelling in dorsal horn, was assessed using the Whole Object Assessment Mode, whereas EGFP+ axon labelling in dorsal horn was assessed using the Object Fragment Assessment Mode. The data output was saved to the user-designated output directory for use in further analysis.

The segmented DRG data was assessed with the Object Manager to confirm the quality of the segmentation, and to define a machine learning classifier to isolate the DRG neuronal nuclei class of objects. Samples of objects across all DRGs in a dataset were selected for inspection and classification into three classes: Feature (neuronal nuclei), Non-Feature (non-neuronal nuclei), Connected (neuronal nuclei connected through the segmentation process). 100 objects across all images were selected randomly at first, to assess the segmentation procedure. Once a high quality segmentation procedure had been confirmed, subsequent rounds of object sampling of 100 objects were biased towards the decision boundary of an interim machine learning classifier, to iteratively bias selection to objects with which the current classifier was most unsure. Using this boosting regime, the machine learning classifier plateauxed its performance in three rounds of object selection (300 objects across all images).

Several passes through the Threshold Manager and Object Manager were required here to define the optimal segmentation of DRG data, to reduce the number of connected objects to low levels, and to define a segmentation procedure which produced a good representation of DRG neuronal objects.

For the DRG datasets, the classifier which consistently performed well was the KStar classifier^36^, an instance-based classifier that uses an entropic distance measure for determining object class.

The machine learning classifier was tested by using an independent test set, created by manually classifying an independent set of objects across all images independent from the original training set objects. This independent test set was classified with the machine learning classifier, and the results comparing the machine learning algorithm with the manual assessments presented in a contingency table.

The segmented axon and synaptic data was assess with the Object Manager plugin to confirm the quality of the segmentation by manual assessment of a random sample of objects across all images. Random samples of 200 objects were assessed for their accuracy in representing the biological object shown in the underlying 8-bit data, and inspected to ensure each represented a single object. No classification was required for these datasets.

### StereoMate Analyser plugin

The StereoMate Analyser plugin implements the stereological assessment of objects, based on the image segmentation protocol developed and tested with the Threshold and Object Manager plugins, and within any user-defined ROIs from the ROI Manager plugin. This plugin uses the output of these previous plugins for input to ensure the correct data and procedures are used for image processing. The plugin runs a non-interactive process to assess all objects in 3D, applying the ROI Di-Sector to objects throughout the image stack. Each image will generate quantitative data for all objects and ROIs, which are output as csv files in the user-designated output directory.

The DRG datasets were processed with the relevant image segmentation procedure and machine learning classifier as previously defined by the Threshold and Object Manager plugins, by selecting the appropriate Object Manager directory as input. No ROIs were defined for whole DRG data, and so for each DRG image stack, the whole image was assessed as one ROI. As the whole region was captured in the stack, the Exclusion Zone XY and Exclusion Zone Z options were left unselected.

The axonal and synaptic datasets were processed with the relevant image segmentation procedure defined previously by the Threshold Manager and Object Manager plugins, by selecting the appropriate Object Manager directory as input. The ROIs previously defined by the ROI Manager plugin were selected as the Input ROI(s). Both the Exclusion Zone XY and Exclusion Zone Z options were selected for pre-synaptic terminal datasets, and these are not selectable for axonal datasets, as these data are analysed as object fragments.

The output of the StereoMate Analyser data was imported into R for subsequent statistical analysis and the generation of graphical plots.

For further details on using the StereoMate platform, please refer to the detiled documentation on the software’s GitHub page.

### Stereological Analysis: Synthetic Datasets

Images of systematically randomly placed objects were generated via an imagej macro by randomly allocating objects to a series of 128×128 pixel blocks. Objects were either squares (16×16, 32×32, 64×64), circles (diameters of 16, 32, 64), or lines of vertical, horizontal or diagonal orientations (4× 16, 32, 64). A sampling region measuring 1280×1280 was taken at random from a larger parent image, to ensure random sampling of the objects. All objects were counted, rejecting any which touched the image edges. For the ROI DiSector, a larger sampling region was used, adding the maximum length of the object under assessment to each dimension. Analysis consisted of counting all objects, but rejecting all objects which contacted the image edge, and also removing any objects which fall completely within the exclusion zone.

### Manual simple and stereological Analysis of DRGs

Simple counts were performed in 3D, and every instance of a DRG neurone identified in a 9 slice (9um) thick Z stack was counted, with no application of a stereological correction.

DRG image stacks were re-sampled for stereological analysis by systematically deriving substacks 20 slices (20um) thick, taken every 10 slices. A depth of 9um were assessed per sample, using the first slice as an initial look-up section, and the remaining 10 slices as an exclusion zone (to allow confirmation if new objects at the bottom of the stack were neuronal nuclei or not, as well as their full reconstruction).

Both simple and stereological counts were performed on systematic random samples across 5 DRGs, using a sampling rate of 1/3 or 1/4 of all substacks. To derive a total neuronal number estimate, the DRG neurone number was multiplied by the sampling denominator.

### Figures and Statistics

FIJI (Fiji Is Just ImageJ) was used for creating all image panels in figures. R and R Studio was used for all graphs and statistical assessments. Figures were arranged using Inkscape.

